# Vision and touch used for catching small balls with a power grip and large balls with a precision grip supports dual visuomotor channel theory

**DOI:** 10.1101/2025.07.24.666538

**Authors:** Amirhossein Mazrouei, Youssef Ekladuce, Hardeep Ryait, Majid Mohajerani, Jenni M. Karl, Ian Q. Whishaw

## Abstract

The dual visuomotor channel theory of grasping posits that distinct neural pathways mediate hand shaping in response to a target’s extrinsic (e.g., location) and intrinsic (e.g., size and shape) properties. To evaluate this theory, we examined grasp behavior in human participants as they caught balls of four diameters (2.5–9 cm) thrown toward them. Hand shaping during catching was compared with that observed during the pickup of stationary balls and the interception of rolling balls. Kinematic measures included digit opposition distance (thumb pad to index finger pad) and prehension span (digit pad to palm distance), obtained using electromagnetic sensors, 3D video capture, and frame-by-frame video analysis. Participants displayed significantly greater hand opening when catching thrown balls than when interacting with static or rolling balls. Nonetheless, the maximum pregrasp aperture (MPA), contact grasp aperture (CGA), and terminal grasp aperture (TGA) scaled proportionally with ball size across all conditions. Ball size further influenced grasp type: small thrown balls were caught with power grips, while larger balls were caught with precision grips. In contrast, precision grips were used consistently when picking up stationary balls or grasping intercepting rolling ones. In the catching condition, grasp type and the trajectory of digit closure were also affected by the location of ball to hand contact. These findings support the dual visuomotor channel theory by demonstrating that anticipatory hand opening reflects target location, whereas grip selection reflects target size. Moreover, the modulation of grasp type and digit closure by tactile contact suggests that somatosensory input may operate within a dual-channel framework analogous to that of vision.

---

Many animal species, including humans, are capable of intercepting moving targets during feeding or play. In humans, the ability to adjust hand movements in response to the spatial and perceptual demands of moving targets emerges early in development (von Hofsten 1983, 1984; Sacrey et al. 2012). Despite this early onset, the visuomotor processes underlying hand shaping toward moving targets and object acquisition are complex (Alderson et al. 1974). Interception studies suggest that movement accuracy is supported by continuous online updating of sensory feedback, allowing for real-time adjustment during task execution (Brenner and Smeets 2018; Slupinski 2018). These findings suggest that while initial movement predictions are important, they are often superseded by moment-to-moment sensory guidance. Previous research on catching has primarily focused on reconciling target motion with the movement of the body, especially in relation to head and eye position or self-motion of the catcher (Cesqui 2016; Höfer 2018; Lacquaniti and Maioli 1989; Savelsbergh et al. 1991, 1993; Laurent et al. 1994; Button et al. 2002; Mazyn et al. 2006; Tijtgat et al. 2010; Bongers et al. 2012). These studies emphasize that successful catching relies on processing both extrinsic (e.g., location, orientation) and intrinsic (e.g., size, shape) object properties in relation to the moving body, particularly the arms and hands. In contrast, hand shaping during catching has received comparatively little attention, particularly when compared to the extensive literature on grasping static objects or intercepting rolling ones (Arbib 1981; Jeannerod 1981, 1999; Jeannerod et al. 1994; Binkofski et al. 1998; Culham and Valyear 2006; Fattori et al. 2010; Cavina-Pratesi et al. 2010a, 2010b, 2018; Karl and Whishaw 2014; Karl et al. 2018; Smeets and Brenner 1999; Vesia et al. 2013; Whishaw et al. 2016).

The present study investigates how the hand shapes and closes to grasp a target in flight. Participants were asked to catch balls of four different diameters thrown toward them. Hand shaping during catching was compared with that observed during grasping of stationary balls and interception of balls rolling across a surface. Based on previous work, we hypothesized that all three conditions — catching, reaching-to-grasp, and reaching-to-intercept — would show common grasp kinematics, reflecting a shared control architecture for hand shaping. Alternatively, we considered that grasp kinematics may vary across conditions due to differing task demands, consistent with the dual visuomotor channel theory (Arbib 1981; Jeannerod 1981; Castiello et al. 1992a; Goodale et al. 1994; Hoff 1993; Hu et al. 2005; Coats et al. 2008; Hoffmann et al. 2010; Begliomini 2014; Karl et al. 2013; Hall et al. 2014). Kinematic variables were measured using electromagnetic trackers, 3D video recording, and frame-by-frame video analysis (Karl et al. 2012, 2013, 2018; Sacrey et al. 2012; Thomas et al. 2014). We quantified maximum pregrasp aperture (MPA), contact grasp aperture (CGA), and terminal grasp aperture (TGA), corresponding respectively to the hand’s maximum opening prior to contact, aperture at first contact, and final aperture during object acquisition. Measurements were expressed in terms of prehension distance (digit pads to palm) and opposition distance (thumb pad to index finger pad), as originally defined by Napier (1956, 1962). Grasp synergies, defined as coordinated digit movement patterns, were analyzed in terms of finger closure sequences and final grasp configuration (Santello et al. 1998; Mason et al. 2001; Lang and Schieber 2004).

For the study, participants were divided into groups for electromagnetic tracking and 3D video capture. However, video recordings were reviewed across all groups to verify and extend kinematic findings and to explore task-related variations in hand shaping and grasp performance.

## Materials

### Participants

A total of 52 right-handed young adults (24 female), aged between 20 and 30, were recruited from the neuroscience subject pools at the University of Lethbridge and Thompson Rivers University. Participants provided informed consent, self-reported no history of neurological, sensory, or motor disorders, had normal or corrected-to-normal vision, and were confirmed as right-handed based on a handedness questionnaire. Ethics approval was granted by the University of Lethbridge, Thompson Rivers University, and University of Alberta Human Subject Research Ethics Committees.

### Electromagnetic monitoring

Hand kinematics were acquired at a 60 Hz sampling rate using a trakSTAR® (Ascension Technology, https://about.ascension.org/our-work/ascension-technologies) system. Electromagnetic sensors were attached to the distal phalanges of the thumb and index finger, as well as on the anterior aspect of the ulnar styloid of the wrist. Spatial coordinates were transmitted with respect to the transmitter, which was fastened atop a pedestal positioned on the floor beside the participant’s right chair leg such that the transmitter and the hand’s start position were vertically aligned. Distinct trials were characterized by a threshold wrist velocity of 5 mm/sec at reach onset and concluded once the target was acquired (grasped, stopped, or caught). Kinematic data capturing the x, y, and z coordinates of the digits were collected to assess hand aperture.

### FreeMocap monitoring

Software used for analysing the videos obtained from three-camera monitoring was FreeMocap (https://github.com/freemocap/freemocap). FreeMocap is open-source software that uses MediaPipe (https://ai.google.dev/edge/mediapipe/solutions/guide) as a base model for 2D pose estimation and then uses triangulation algorithms to perform 3D pose estimation. The videos were synchronized for FreeMocap using Adobe Premiere Pro (https://www.adobe.com/ca/).

### Video recording

In the electromagnetic monitoring experiments (see below), participants were monitored with two VIXIA HF G70 Canon video camcorders, set at 30 Hz fps rate and 1/1000 shutter speed, with one camera capturing a frontal view and one camera capturing a lateral view. Three GoPro 11 cameras (gopro.com) were used in the FreeMocap recording experiments (see below) to capture frontal and side views, recording at 120 frames per second with a shutter speed of 1/960 seconds.

### Balls

Four soft balls with diameters of 2.5, 3.5, 6, and 9 cm were used. The 6 cm diameter ball was a tennis ball (Wilson Open Extra Duty), while the others were made of soft rubber.

### Digit designation

The digits were designated numerically from 1 to 5, with 1 representing the thumb and 2 through 5 representing the fingers, starting with the index finger and ending with the pinky.

### Grasp scoring

Grasp scoring was made from the electromagnetic measures that indicated kinematic events, and these were processed offline using the electromagnetic readout and by stepping frame-by-frame through the video. Measures made were:

*(a) Maximum pregrasp aperture (MPA).* MPA was defined as the largest opposition distance between the thumb and index finger occurring after reach onset and prior to the grasp.
*(b) Contact grasp aperture (CGA).* The contact grasp aperture reference was determined by inspection of the video record for the point that a ball first touched the hand and then reading the kinematic record to obtain that value.
*(c) Terminal grasp aperture (TGA).* The terminal grasp aperture was the minimal distance between the thumb and index finger occurring after contact aperture. This represented finger closing on a ball as confirmed by visual inspection of the video record.
*(d) Precision grasps.* Precision grasps (Napier 1956, 1962) are those in which a ball is held between one or more terminal digit pads and the terminal pad of the first digit (thumb). Grip type was determined by frame-by-frame inspection of video made from either frontal or lateral perspectives and the scores were confirmed from the kinematic measures.
*(e) Power grasps*. Power grasps (Napier 1956, 1962) were those in which an object was held by one or more fingers against the palm. Some power grasps had counter pressure applied by the thumb, in which the thumb is positioned over and applies pressure against the second digit. Other power grasps had the thumb extended away from the grasping fingers. Grip type was determined by frame-by-frame inspection of video made from either frontal or lateral perspectives and the scores were confirmed from the kinematic measures.
*(f) First hand contact*. The first hand contact with a ball was rated on a three-point scale: 0 - first contact was with a digit pad, 0.5 - first contact was with the ventral surface of a digit, 1 - first contact was with the palm.
*(g) Palm contact*. Palm contact by a ball was identified through frame-by-frame inspection of the video recordings. The location where the ball first touched the palm in the catching experiments was scored corresponding to palm sections continuous with one of the five digits, in which 1 represented the palm region aligned with digit 1 (thumb) and 2, 3, 4, and 5 represented the region of the palm aligned with the other digits respectively.
*(h) Reach*. The reach was reconstructed by using the x-y coordinates from the automated video tracker on the digit tip of the 5th digit, the wrist, elbow, and shoulder in order to generate a time-distance-velocity curve and an x-y trajectory for a catch. The measurements were taken from movement initiation until the moment the grasp was completed.
*(i) Chase catch*. A chase catch was a catch in which there was a quick acceleration of the hand after ball contact that was associated with closing the digits to grasp. A chase catch has features that might be associated with catching a flying insect. A catch was scored as being a chase catch or not.
*(j) Stop catch*. A stop catch was one in which the hand both stopped an airborne ball and closed the digits to grasp it, with minimal other hand movement. A stop catch was scored as being a stop catch or not.
*(k) Finger use in precision grasps*. The digit that was in direct opposition to digit 1 (thumb) for a precision grip for grasping a ball was defined as the digit making the main contribution to a precision grasp. The grasping patterns were scored from “2 to 5”, in relation to terminal pads of the four digits. If a ball was held with a power grip, the grip type was given the designation of “1” (Figure 1, right).
*(l) Palm contact location*. Palm contact by a ball was determined by frame-by-frame inspection of the video record. The location of the ball when it first contacted the palm in the catching experiments was scored in relation to palm sections that were contiguous with one of the five digits and numbered 1-5.

**Figure 1.**
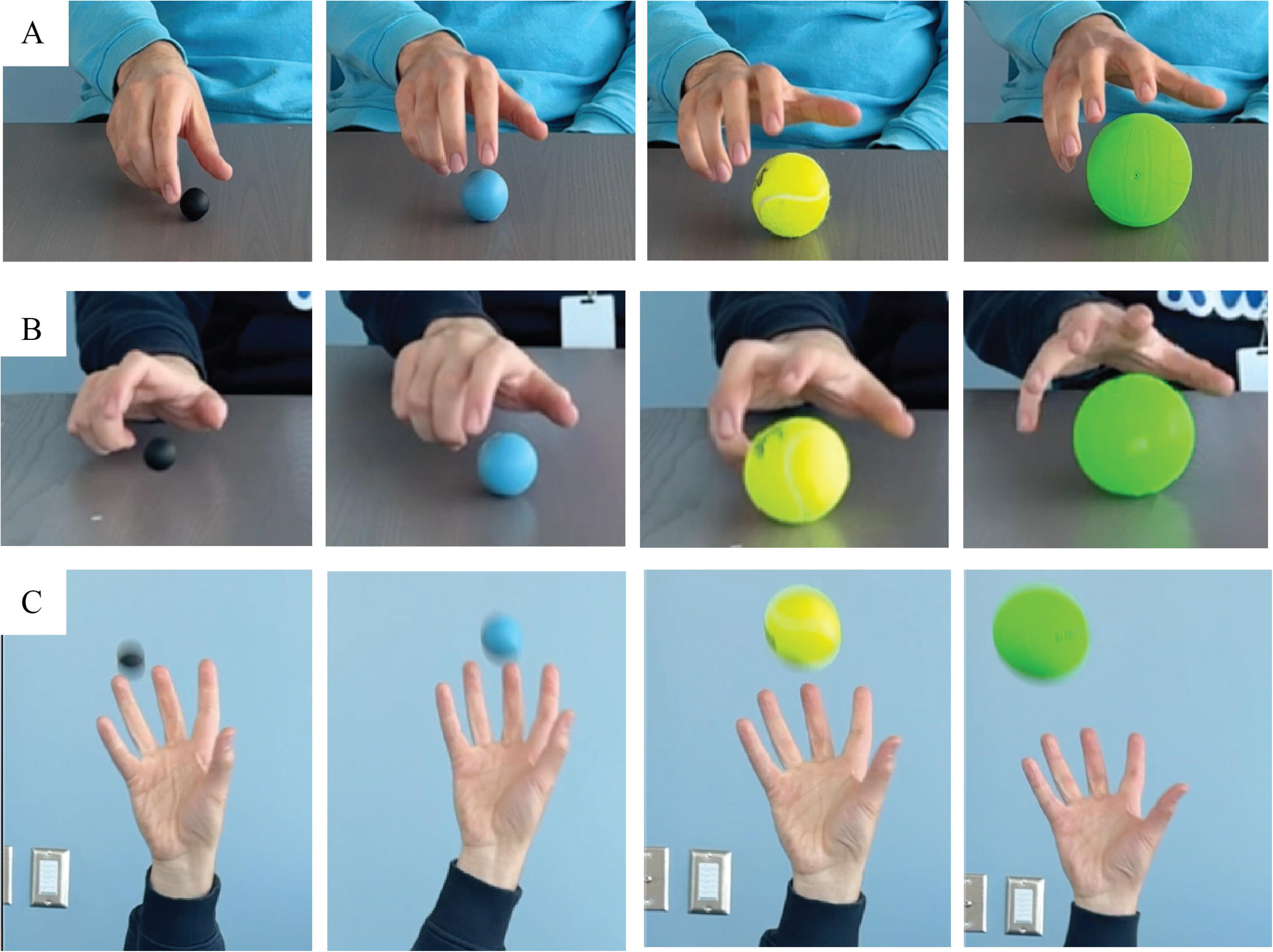
Hand shaping at maximum pregrasp aperture (MPA) in three grasp tasks. Hand shaping at MPA, as estimated for opposition distance between the distal thumb and first finger, four 4 balls of different size in tasks of: A. reach-to-lift, B. reach-to-intercept a rolling ball, and C. reach-to-catch a thrown ball. MPA increased as a function of ball size in all grasp tasks. In addition, the MPA of the hand was larger for catching balls of all sizes in that it featured a nearly fully opened hand, although opposition distance between the thumb and first finger did increase with ball size.

### Hand measurements from FreeMocap monitoring

The following measures of hand shaping during catching, derived from 3D video, were included: opposition distance, prehension distance, and hand rotation.

*(a) Opposition distance*. Opposition distance (Napier 1956, 1962) represents the relationship between digit 1 (thumb) pad and the distal pads of the other digits. Opposition distance was measured as: With *T_tip_* denoting the 3D coordinates of the thumb distal pad, and 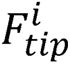 represent the 3D coordinates of the *i*-th distal finger pad, where *i* ∈ {index, middle, ring, pinky}. The distance (di) was computed as:

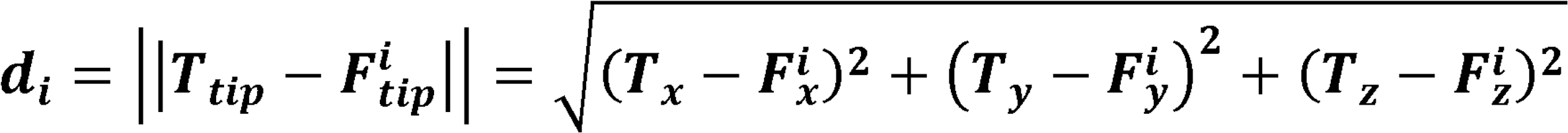

*(b) Prehension distance.* Prehension distance (Napier 1956, 1962) represents the distance from the distal digit pads to the plane of the palm at the palm surface. Prehension distance was measured as: The palm plane was defined using three anatomical landmarks, index finger MCP (metacarpophalangeal joint) root (*I_mcp_*), pinky finger MCP root (*P_mcp_*), and wrist center (*W*). The plane equation *ax* + *by* + *cz* + *d* = 0 was derived from the three points *I_mcp_*, *P_mcp_* and *W*. From the vectors in the plane 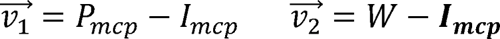 the normal vector 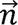 to the plane was 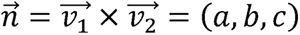 and was solve for d using *I_mcp_* with 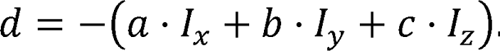. Prehension distance was then calculated as the distance (*D_i_*) from a fingertip 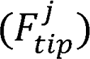 (where *j* ∈ {thumb, index, middle, ring, pinky}) to the plane as:

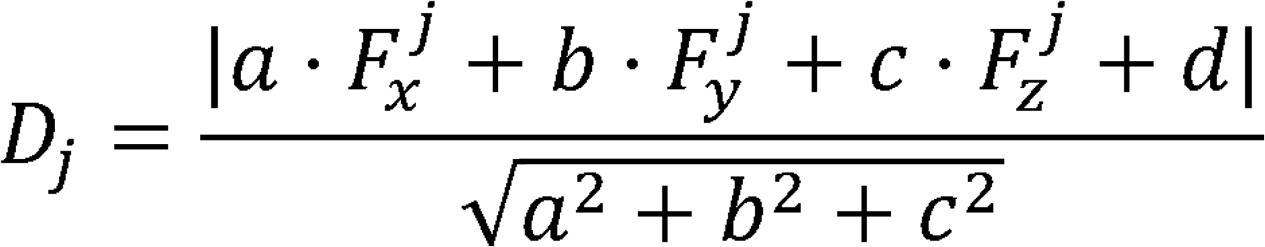

## Procedure

Before starting the experiment, participants were briefed on the task and completed a handedness questionnaire (Bryden, 1977) to confirm their preference for using the right hand when picking up objects and catching a ball.

### Task comparisons

Participants were fitted with electromagnetic sensors (6 male, 14 female) were seated in a comfortable upright position in an armless chair with their feet flat on the floor and their hands resting in the start position open on their lap. They completed three tasks in which they grasped balls of four different sizes 6 times in each task.

(a) *Reach-to-grasp a stationary ball task*. In this task, a stationary ball was placed on the table at a comfortable, predetermined arm’s-length distance. Participants were instructed to reach out, grasp, and lift the ball without any specific instructions on how to perform the grasp.
(b) *Reach-to-grasp a rolling ball task*. In the rolling ball task, a ball was rolled by the experimenter diagonally across the table toward the participant and the participants were asked to intercept the ball by grasping it.
(c) *Reach-to-catch a thrown ball task*. For this task, the experimenter stood opposite to the right hand side of the participant and using an underhand throw, threw the ballt o a point above the participant’s right shoulder , allowing the participant to catch the ball with an overhand catch.

### Catching balls at different locations

Participants (5 male, 5 female) were required to catch each of the four different sized balls six times in each of three different locations while standing. Locations were:

(d) *Overhand-catch*. In the overhand-catch task, the ball was thrown underhand to just above the participant’s right shoulder, prompting them to catch it with an overhand grasp..
(e). *Chest-catch*. In chest-catch task, an underhand thrown ball was aimed to the right side of the participant at chest level.
(a) *Underhand*-catch. In underhand-catch task, A ball was thrown underhand to the midpoint of the body on the participant’s right-hand side. Participants were asked to catch the ball with an underhanded catch.

### 3D filming task

Participants (10 male, 10 female), filmed in 3D, were seated and completed six catching trials with each of the four different-sized balls. The experimenter stood opposed to the right hand of the participant and using an underhand throw, directed the ball to a point above the right shoulder of the participant, so that a participant caught the ball with an overhand catch.

## Statistical Analysis

Statistical analyses were performed using the software IBM SPSS Statistics (v28.0.1.1). The data are presented as mean ± standard error of mean (mean ± sem). For statistical comparisons, a mixed model ANOVA was used for within and between subject measures. A p-value of <.05 was defined as statistically significant. Power was measured by eta-squared (*n*^2^), a descriptive measure of the strength of association between independent and dependent variables. Power analysis was used to confirm that the minimum sample size observed for a conclusion was adequate. Bonferroni tests were used for follow-up comparisons, also with a significance level of p < .05. Pearson product-moment correlations were used for data fits and Pearson product correlations were expressed as r. For principal component analysis (Everitt al. 2011) of the sequence of digit closing to grasp, the data was first transformed into a binary structure, where each column represents whether a specific digit was closed at a particular order (e.g., ’order1, digit indicates if digit 1 was closed at order 1). PCA was then applied for dimensionality reduction. The optimal number of clusters was determined using the Elbow method on the PCA-transformed data. K-means clustering was performed with the selected number of clusters, and cluster characteristics were analyzed by examining the mean values of the original binary variables within each cluster.

## Results

### Catching features larger hand apertures than lift or intercept

Figure 1 illustrates representative hand shapes at MPA, as estimated for opposition distance between the pads of digit 1 (thumb) digit 2 (index finger). Apertures for the four 4 different sized balls are shown for the three tasks tasks: (A) reach-to-lift, (B) reach-to-intercept a rolling ball, and (C) reach-to-catch a thrown ball. MPA catching balls of all sizes the hand featured a nearly fully opened hand. Nevertheless, MPA was proportional to ball size, as indicated by the following statistical results.

Figure 2 illustrates differences (mean±sem) in hand shaping in the three task as represented by opposition aperture between the digit 1 (thumb) distal pad and the digit 2 pad. The reach-to-catch task features a near fully opened hand at MPA (Figure 1A-left) and at CGA (Figure 1B-left) for balls of all sizes. Measures from electromagnetic monitoring of opposition aperture between the thumb and the first finger indicated that hand aperture increased in size in the order of reach-to-grasp < reach-to-intercept < reach-to-catch. Apertures at TGA were similar for all tasks (Figure 1C-left) The larger MPA and CGA associated with catching was not influenced by whether catches were made overhead, at chest level, or underhanded (Figure 1ABC-right). These results were confirmed by the following statistical analyses.

**Figure 2.**
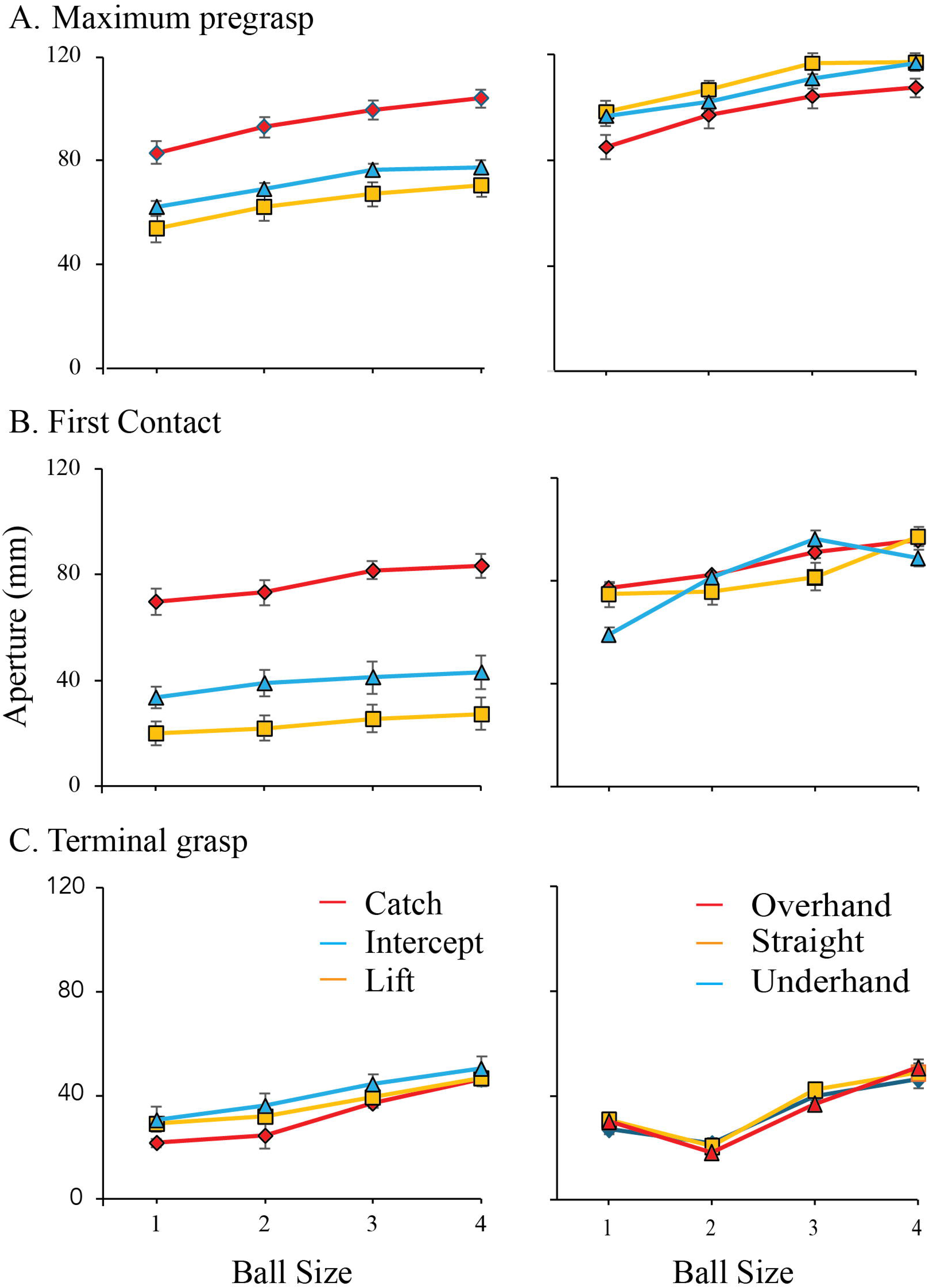
Thumb and first finger opposition associated with catching different sized balls. A-left. Maximum pregrasp aperture (MPA), as measured as opposition distance between the thumb and first finger, associated with catching balls for four different sizes in tasks of reach-to-lift a static ball, reach-to-intercept a rolling ball, and reach-to-catch a ball thrown above the shoulder. B-left. The first contact aperture (FCA) between the thumb and first finger, occurring with the first contact of the hand with a ball, for the different sized balls in the three tasks. C-left. The terminal grasp aperture (TGA), associated with final grasping the different sized balls in the three tasks. Right. Measures of thumb to first finger opposition, for catching balls of four different sizes, above the head, at chest level, and underhanded. Notes: (1) hand aperture increases with ball size, (2) MPA and FCA are larger in the catching tasks (the apertures associated with catching the smallest ball were larger than the apertures associated with lifting or intercepting the largest ball), (3) the hand aperture associated with catching was did not change as a function of catching location.

*Maximum pregrasp aperture (MPA)*. Figure 2A (left) gives a summary of the MPA, made by the thumb and the index finger, for the three reach tasks with four different sized balls. The ANOVA revealed a significant effect of Task, F(2, 38) = 42.662, *p <0.001*, *η^2^* = 0.692, with the MPA aperture for reach-to-catch > reach-to-intercept > reach-to-grasp (follow-up Bonferroni tests p < 0.001). There was also a significant effect of ball size, Size: F(3, 57) = 112.636, *p < 0.000*, *η^2^*= 0.856, with MPA increasing in size in relation to ball size (ball 1 < ball 2 < ball 3 < ball 4, p<0.001). There was no significant interaction between Task and Ball Size, F(6, 114) = 1.130, *p = 0.56*, *η^2^* = 0.431.

*Contact grasp aperture (CGA). (A)*. Figure 2B (left) gives a summary of the CGA. There was a significant effect of Task: F(2, 38) = 37.093, *p = .001, η^2^* = 0.66, with reach-to-catch > reach-to-intercept > reach to grasp (p < 0.001). There was also a significant effect of ball size, Size, F(3, 57) = 13.575, *p* < *.001, η^2^* = 0.417, with FCA increasing in size in relation to ball size (ball 1 < ball 2 < ball 3 < ball 4, p < 0.001). There was no significant interaction between Task and Ball Size, F(6, 114) = 1.436, *p = 0.207, η^2^*= 0.070.

*Terminal grasp aperture (TGA)*. Figure 1C (left) gives a summary of TGA. There was a significant effect of Task, F(2, 38) = 11.620, *p = .000, η^2^* = 0.379, with TGA aperture measures giving aperture differences of reach-to-catch < reach-to-intercept = reach-to-grasp (p<0.001). There was also a significant effect of ball size, Size, F(3, 57) = 46.282, *p = .000*, *η^2^* = 0.709, with TGA increasing in size in relation to ball size (ball 1 < ball 2 < ball 3 < ball 4, p<0.001). There was no significant interaction between Task and Ball Size, F(6, 114) = 1.041, *p = .403*, *η^2^* = 0.052.

### Open hand aperture for catching at all locations

Figure 2 (right) gives a summary of aperture measures as participants made catches of the four different balls thrown above the shoulder, at chest level, or at waist level. Measures of aperture between the thumb and the index finger indicated that that similar aperture occurred at each catch location, although the overhead aperture was slightly smaller than were chest and waist level apertures. These results are confirmed by the following statistical analyses.

*Maximum pregrasp aperture*. Figure 2A (right) summarizes MPA when catching at different locations. The ANOVA for MPA was significant, Task: F(2, 22) = 7.661, *p = .003*, *η^2^* = 0.411, with comparative grasp aperture measures giving straight = underhand, > overhead (p<0.001). There was an effect of ball Size, F(3, 33) = 61.735, *p = .000*, *η^2^* = 0.849, and aperture was largest for the largest balls, ball 1 < ball 2 < ball 3 < ball 4, p< 0.05 for all reaches. The interaction between Task and Size was not significant, F(6, 66) = 0.926, *p = 0.482*, *η^2^* = 0.078.

*Contact grasp aperture*. Figure 2B (right) gives a summary of CGA when the catch occurred at different locations. There was no effect of catch location on hand aperture at first contact with a ball, Task, F(2, 22) = 0.185, *p = .832, η^2^* = 0.017, but there was a significant effect of ball size, Size, F(3, 33) = 4.750, *p = .007, η^2^* = 0.302, with ball 1 < ball 2 < ball 3 < ball 4, p< 0.05. The interaction between Task and Size, F(6, 66) = 1.173, *p = 0.331, η^2^* = 0.096, was not significant.

*Terminal grasp aperture*. Figure 2C (right) gives a summary of TGA when catching at different locations. For TGA, the was no significant effect of task, F(2, 22) 0.780, *p = .471*, *η^2^* = 0.066. There was an effect of ball Size, (3, 33) = 61.023, *p = .000*, *η^2^* = 0.847, ball 1 > ball 2 < ball 3 < ball 4, p<0.05. The larger hand aperture associated with the smallest ball than the second smallest ball in Figure 2C (right) likely related to differences in the type of grasp and this is discussed below. The interaction between Task and Size, F(6, 66) = 1.446, *p = .211*, *η^2^* = 0.116, was not significant.

### Opposition and prehension measures show all digits open for catching

Figure 3 illustrates representative velocity kinematic curves of all digits from 3D videorecords for the smallest ball and the largest ball. The curves illustrate that all digits open in parallel in association with hand opening to catch. Figure 3A shows opposition distance, the distance between the distal thumb pad and the distal pads of each finger. Figure 3B shows prehension distance, the distance between the distal digit pads and the palm.

**Figure 3.**
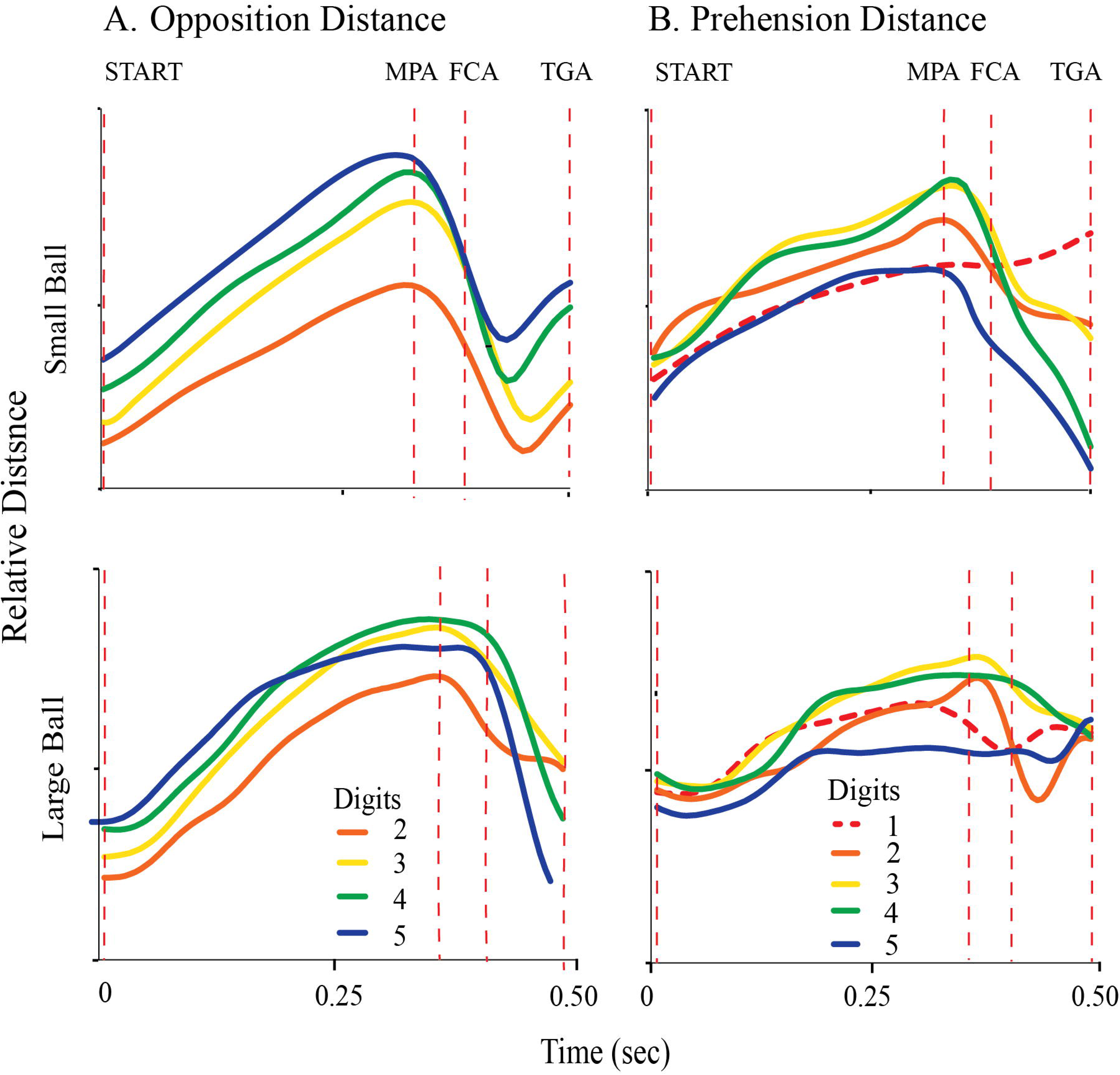
Kinematics of finger shaping associated with catching. Left, opposition distance, the distance from the terminal pad of the thumb to the terminal pads of each finger; Right, prehension distance, the distance from the terminal pads of each digit to the palm. The vertical dotted lines indicate; START, the point that the hand is lifted with the fingers lightly flexed in a collect posture, Maximum Pregrasp Aperture (MPA), the largest opposition distance between the thumb and each finger, First Contact Aperture (FCA) and Terminal Grasp Aperture (TGA). Top a representative catch of a small ball, Bottom, a representative catch of the larger tennis ball. Note: (1) relative synchrony of finger opening and closing, (2) the variation in the kinematic curves following MPA is likely related variations in catching strategies.

Figure 4 summarizes, (A) opposition distance and (B) prehension distance, for each digit at MPA, CGA and TGA, associated with catching the balls of four different sizes. Both measures reveal that the aperture of the hand was closing both prior to contact with the ball and after contact. Opposition measures of the distance between the distal pad of the thumb and the distal pad of each of the 4 digits gave a significant effect of the three aperture measures, F(2,18)=92, p<0.001, *n*^2^= 0.96, power = 1, and follow-up tests indicated that MGA > FCA > TGA, p<0.001. Prehension measures between the distal pads of the finger and the palm were also significant, F(2,18)=810, p<0.001, *n*^2^= 0.96, power = 1, and follow-up tests indicated that MGA > FCA > TGA, p<0.001.

**Figure 4.**
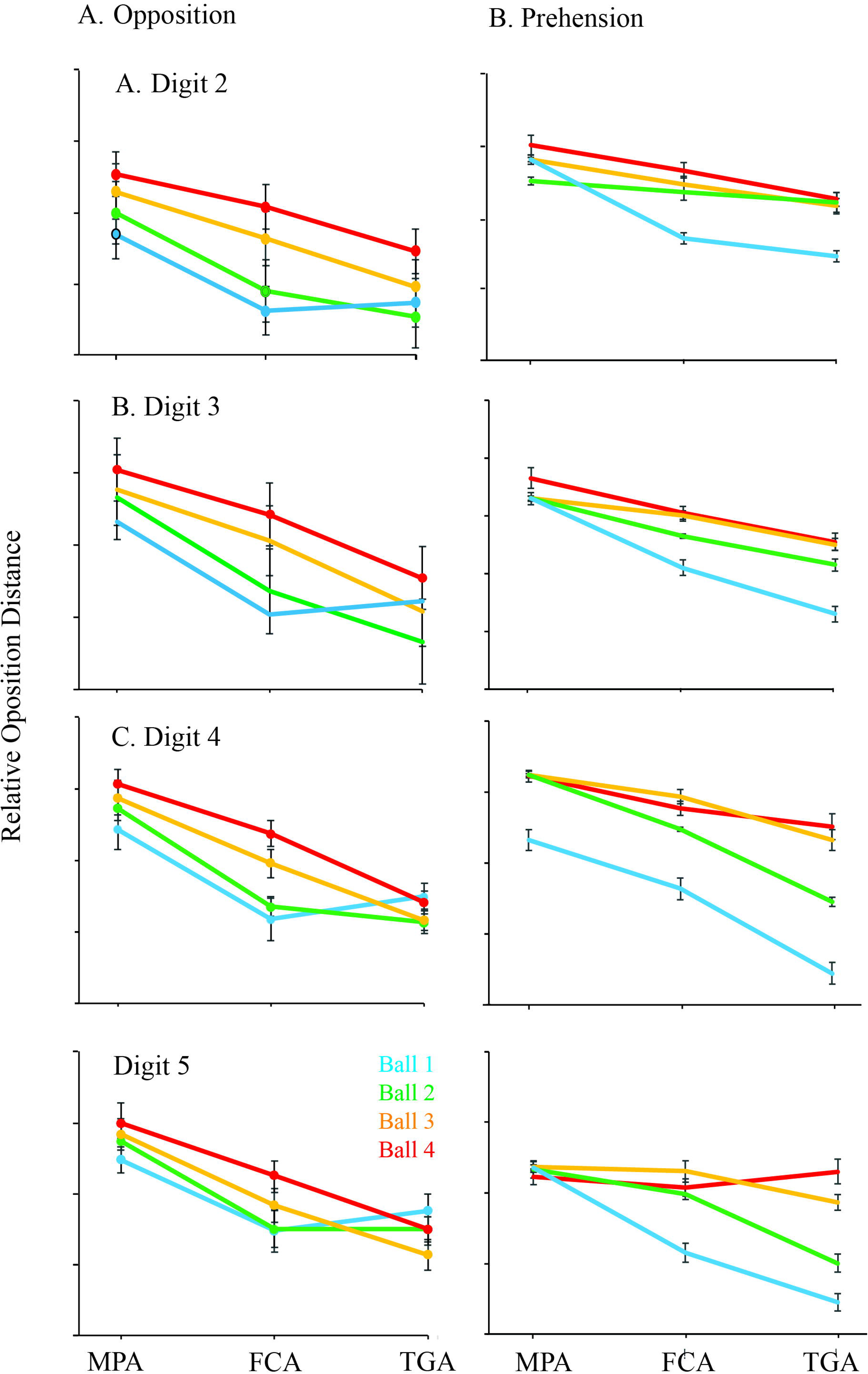
Opposition and Prehension reveal digit closing before and after first contact. A. Opposition aperture, distance between the terminal thumb pad and the terminal pad of each finger. B. Prehension aperture, distance from digit pad to palm. Measures are made at Maximum Pregrasp Aperture (MPA), First Contact Aperture (FCA) and Terminal Grasp Aperture (TGA). The aperture of each digit for catches for balls of four different sizes are represented. Note: (1) the relative synchrony of digit opening and closing, (2) apertures are larger for larger balls, (3) digit closing occurs before hand contact with a ball and after hand contact.

Despite the synchrony in digit closing to catch there were differences in the aperture of each of the digits in relation to ball size (opposition, F(3,27) = 124.1, P < 0.001; prehension F(3,27) = 32.0, p<0.001). Follow-up tests showed that hand aperture for Ball 4 > Ball 3, > Ball 2, > Ball 1, p < 0.001 for both opposition and prehension.

There were significant interactions between ball size, digits, and aperture measurements. These largely reflected the variations in grip applied to the balls of different size after first contact between the hand and a ball (see below). Significant interactions included: ball size by digit (opposition F(9,81) = 35.8, p < 0.001; prehension F(12,108) = 5.6, p < 0.001); digits by aperture measure (opposition F(6,54) = 15.3, p<0.001; prehension F(8,72) = 21.9, p<0.001) and measures by ball size by digits (opposition F(18,162)=13, p<0.001; prehension F(24,216)=11.39, p<.001).

### First contact made with the palm for catching

Two measures were made to determine which part of the hand made first contact with the ball in the different tasks. One measure examined whether the terminal digit pads made first contact and the other measure examined hand orientation to the balls at first contact.

### Digit vs palm contact

Figure 5 summarizes results that show that the digit tips made first contact with the small stationary balls. Palm contact was more likely as the balls became larger or were intercepted or caught. Figure 5A is an example of the use of the distal thumb pad and distal finger pads of the first two fingers to make first contact with the smallest stationary ball. Figure 5B illustrates that first contact with the tennis ball is made with the palm and the ventral surface of the thumb and the first two fingers. Figure 5C gives a summary of the results obtained from all participants in the reach-to-grasp, reach-to-intercept, and reach-to-catch tasks. Statistical comparisons of whether the distal digit pads, the ventral surface of the digits, or the palm made first contact gave a significant effect of task, (F2,38) = 12.4, p<0.001. There was also a significant effect of ball size, F(3,57) = 59.2, p<0.001, *n^2^* = 0.8, power = 1, and a significant interaction of task by ball size, F(6,114) = 10.8, p<0.001, *n^2^*= 0.4, power = 1. Follow-up tests on contact scores for both task and ball size gave the effect of reach-to-grasp < reach-to-intercept = reach-to-catch, with the lower values indicating digit tip first contact. In summary, the participants were more likely to make digit contact with small stationary balls and to use the palm to contact with larger balls moving balls.

**Figure 5.**
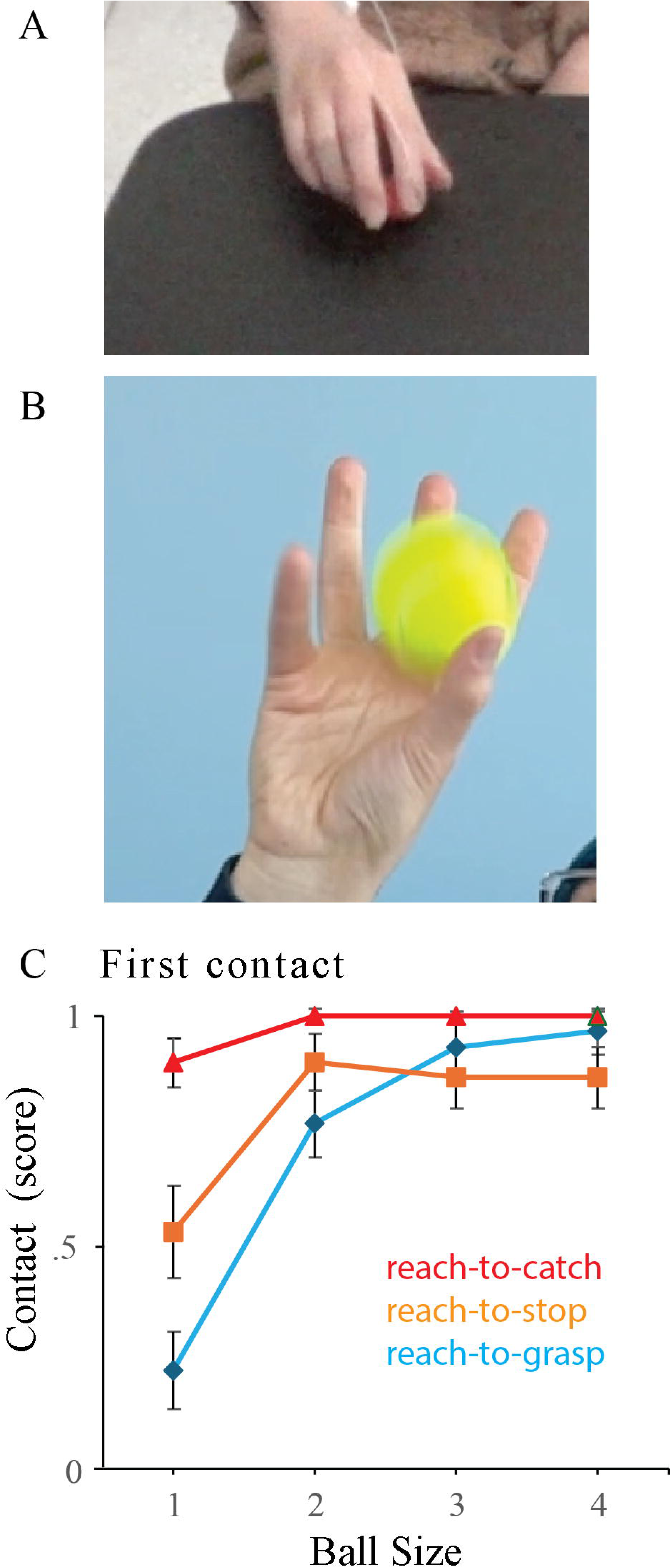
Kinematic representation of a power and precision catch. Illustration and kinematic curves associated with: A. A power catch, in which a small ball is grasped between the closed fingers and the palm with no involvement of the thumb. B. a precision catch, in which a tennis ball is held between the distal pads of the thumb and the distal pads of a number of the fingers. The insert bar graph in B shows the incidence of stop-catches in which the hand is relatively still or moves slightly backward at ball contact. Chase-catches, in which the hand moves forward during the grasp were more common with small ball catches. The right forelimb was digitized at the tip of index finger, the wrist, the elbow and the shoulder.

### Hand alignment for catches

Figure 6 gives a summary of the orientation of the hand to a target for each ball size in each task at first contact. To measure orientation of the hand to a target ball, the palm was divided into 5 vertical segments from the radial to ulnar side of the hand, each of which was aligned with a digit (insert Figure 6 top). Upon first contact with the ball, a score of 1-5 was given to the alignment of the center of the ball and the hand. For the smallest ball in all tasks, the average alignment to the target was toward the radial side of the hand. For larger balls in all tasks, the average alignment was toward the center of the hand. This summary was confirmed by a significant effect of task F(2,38) = 2.5, p=0.10, η^2^ = 0.12 and a significant effect of ball size, F(3,57) = 3.64, p=0.2, η^2^= 0.2, and no significant interaction of task by ball, F(6, 114) = 0.7, η^2^ = 0.3.

**Figure 6.**
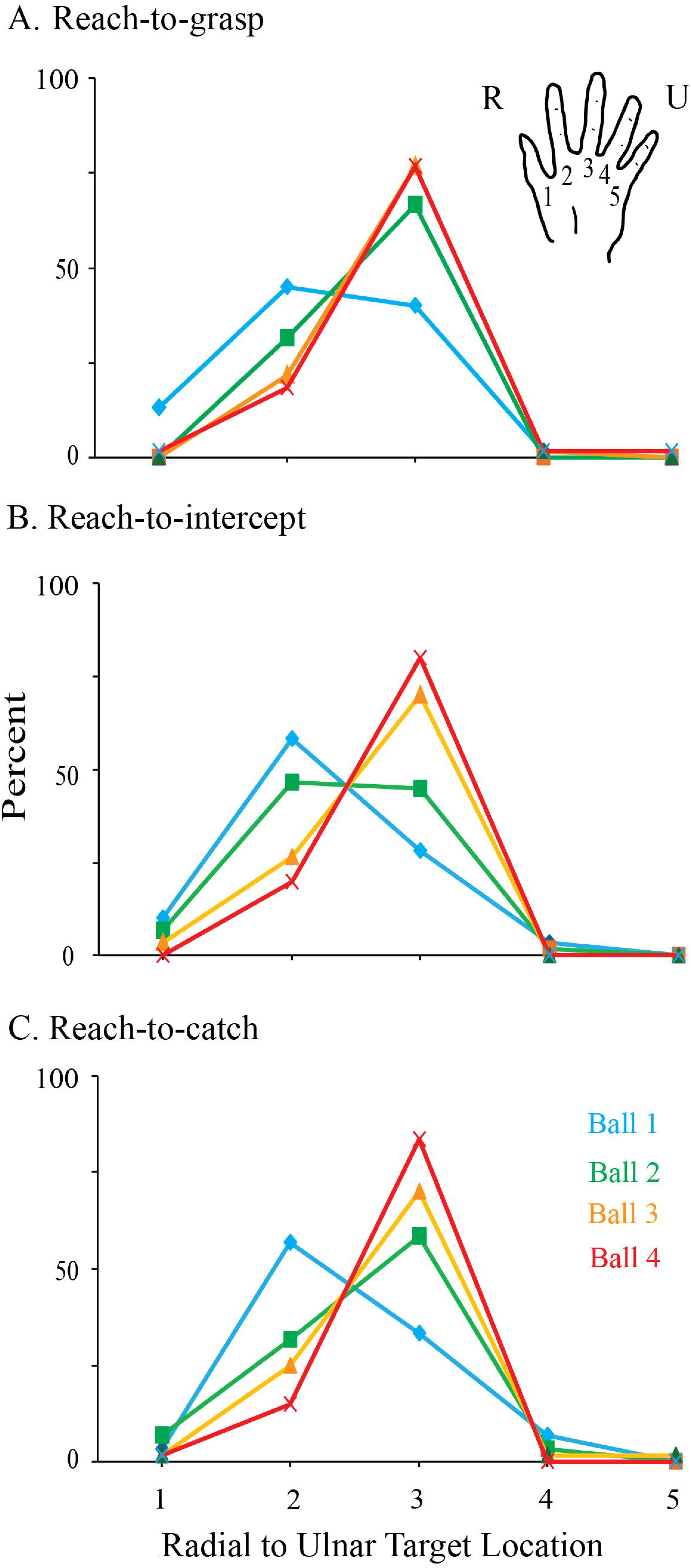
Digit-tip contact with small stationary balls but ventral surface hand contact with large and moving balls. A. A participant lifts a stationary smallest ball using the distal pads of the thumb and first two fingers. B. A participant catches the tennis ball using the lower palm and the ventral surface of the thumb and the first two digits. C. Summary of the location of the first contact between a ball and the hand as a function of ball size and task. Note: low scores indicate first contact with the distal digit pads and high scores indicate first contact with the ventral surface of the digits and/or the palm. Note: A small stationary ball was likely to contacted first by the distal digit pads whereas as large and moving ball were likely to be contacted by the ventral surface of the digits and/or the palm.

## Power grips used to catch small balls and precision grips used to catch large balls

Two grip types were made for catching the balls. Figure 7A (Video 1) gives examples of kinematic velocity profiles for a power grip used to catch the smallest ball and Figure 7B (Video 2) gives the profile for a precision grip used to catch the largest ball. The tip of digit 1, the wrist, the elbow, and the shoulder were digitized for the two grips obtained from the same participant. The velocity profiles indicate that both types of catches are initiated by movement of the lower arm followed by movement of the upper arm, which bring the hand to MPA. The two grasps can differ thereafter, however.

**Figure 7.**
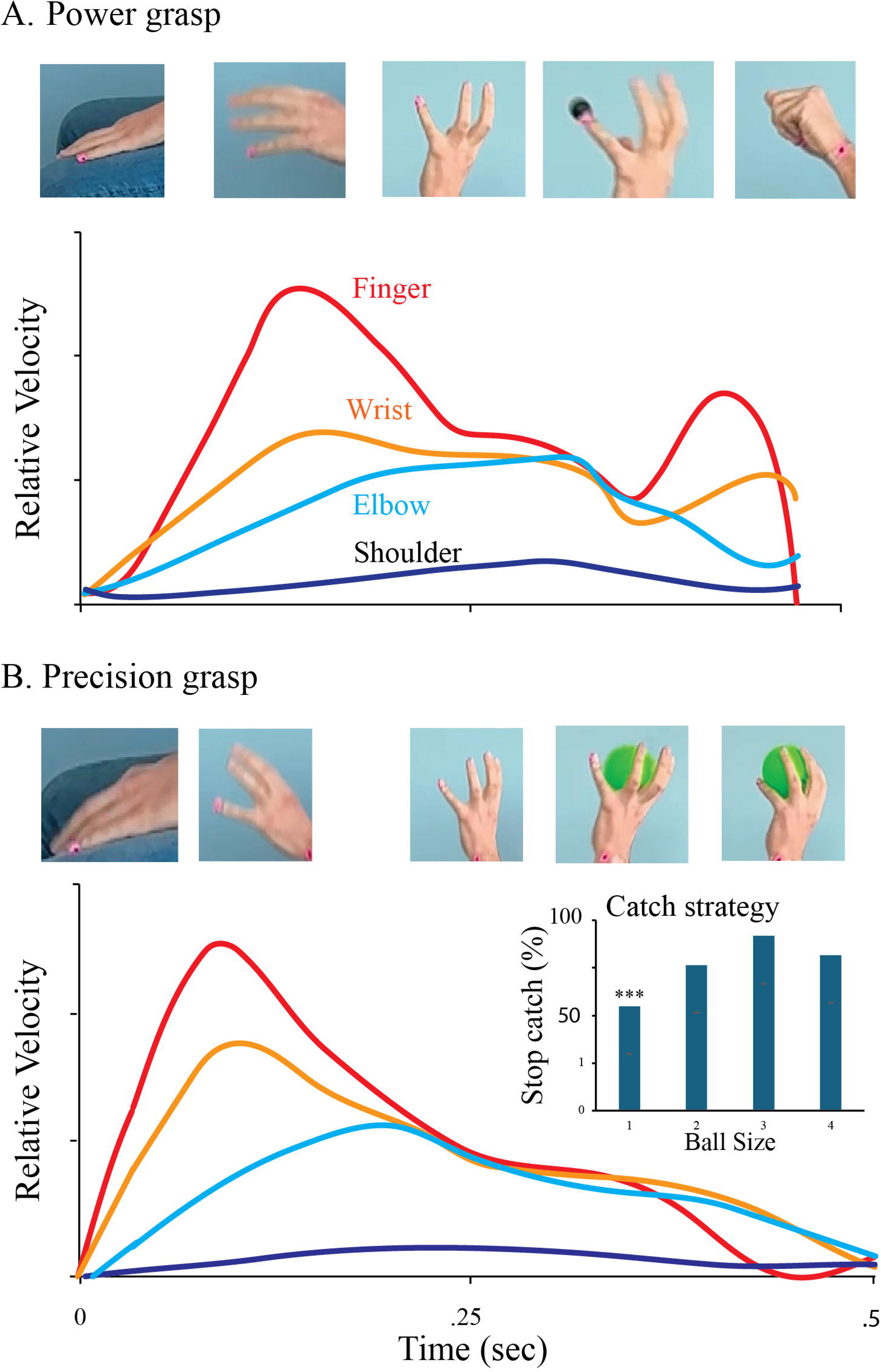
Hand target for grasping. Mediolateral location of first ball contact with the hand as measured in relation to the digits (digit 1 to 5). A. Reach-to-grasp, B. Reach-to-intercept, C. Reach-to-catch. Note: the smallest ball was contacted toward the radial side of the hand whereas larger balls were more likely to be contacted with the midpoint (digit 3) of the hand.

We observed that when the smaller balls struck the palm of the hand they frequently bounced. Consequently, the hand made a rapid forward movement to assist the catch, illustrated by the increase in velocity of the wrist and digit at the end of the catch in Figure 7A. We termed this type of catch a chase catch. Otherwise, the hand retracts somewhat, with a movement of the lower arm. We have termed this type of catch a stop catch. The insert in Figure 7B shows that chase catches were used more frequently for smaller balls and stop catches were used most frequently when catching the larger balls, F(3,57) = 12.9, p>0.001, *n^2^*=0.4, power = 1.

### Features of power and precision grips

A power grip involved closing the fingers to trap a ball between the fingers and the palm, with the thumb usually extending away or sometimes flexing against the outside of the fingers. A precision grip involved grasping a ball between the external pad of digit 1 (thumb) and the external digit pads of one or more of the other digits. Counts of the incidence of the use of the grip types obtained from the first three catching trials from each of 50 participants indicated that power grips were used for 96, 56, 2 and 0 percent of each of the ball sizes. Thus, power grips were mainly used to catch the small balls and precision grips were used to catch the large balls.

We assumed that the main pressure used for holding a ball was exerted through the diagonal midpoint of a ball. Thus, for power grips a ball was held between the digits and palm and for precision grips a ball was held by opposition between digit 1 (thumb) and one of the other digits. Figure 8 gives a summary of estimates of the main oppositions used for purchasing the different sized balls in reach-to-lift, reach-to-intercept, and reach-to-catch tasks. The white bars indicate power grips and the colored bars designate oppositions between digit 1 (thumb) and each of the other digits. The main finding was that for precision grips, as ball size increased, opposition between digit 1 (thumb) and the other digits shifted in a radial to ulnar direction (index to pinky). With dependent variables of power grip and opposition between the thumb and each of digits 2-5, the radial to ulnar shift with ball size gave a significant effect of Task, F(2,36)=6.46, p=0.004, η^2^ = 0.39, power = 0.97; ball size, F(3,54) = 49.8, p<0.001, η^2^ = 0.74, power = 1.0, and no significant interaction between task and ball size, F(6,108) = 1.76, p> 0.05. The task effect was mainly related to the increased use of the power grip for catching vs lifting and intercepting.

**Figure 8.**
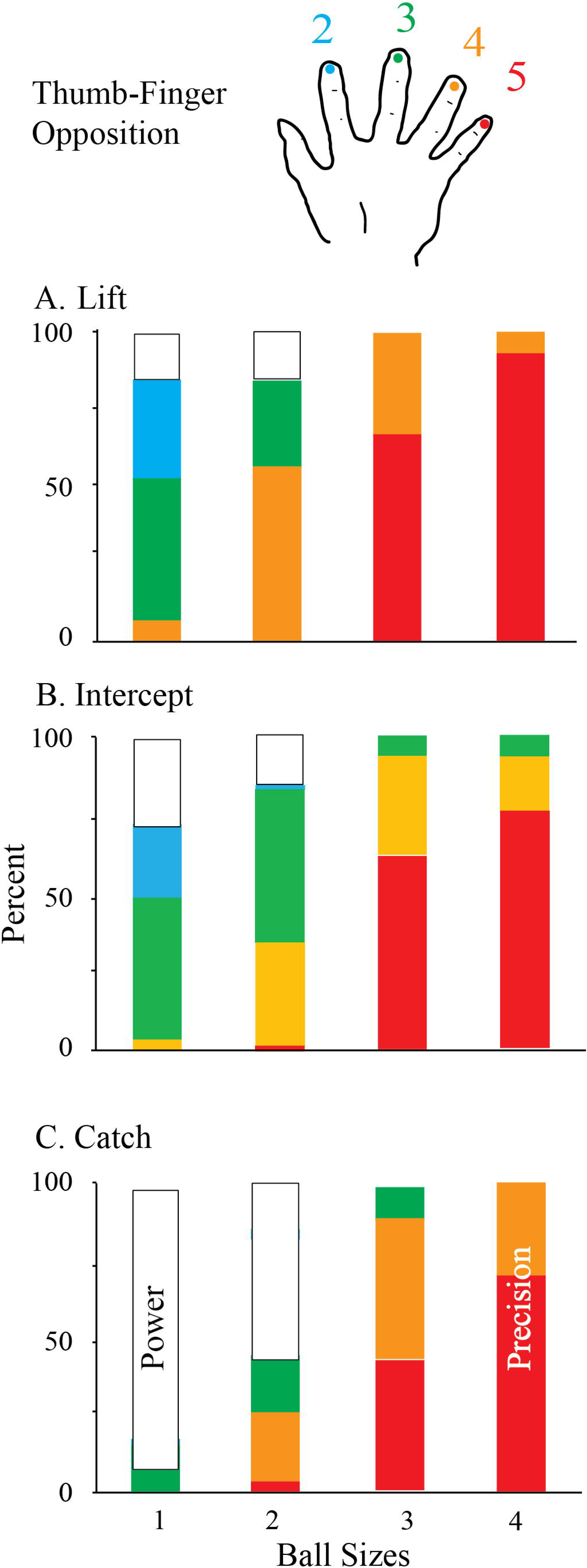
Digit use in the grasp. A. Reach-to-lift, B. Reach-to-grasp, C. Reach-to-catch. The open bars represent power grasps, in which a ball was grasped between the fingers and the palm. The color bars represent the opposition between the distal pad of the thumb and the finger diagonally opposite the thumb (color coding to indicate fingers is indicated by the cartoon of the hand). Note: digit 1, the thumb, is not used for power grasp and the opposition between the thumb and fingers for precision grasps shifts toward the ulnar fingers as ball size increases.

### Grasp synergies influenced by touch

The order of finger closing to make a grasp was observed from a frontal perspective to examine variations in the pattern or synergies with which the distal digit pads closed. the measures were made using catches of the smallest ball, which mainly featured a power grip, and the tennis ball which mainly featured a precision grip. Figure 9 gives a matrix showing the order (as described numerically and by heat map color) of finger closing for power grips and for precision grips.

**Figure 9.**
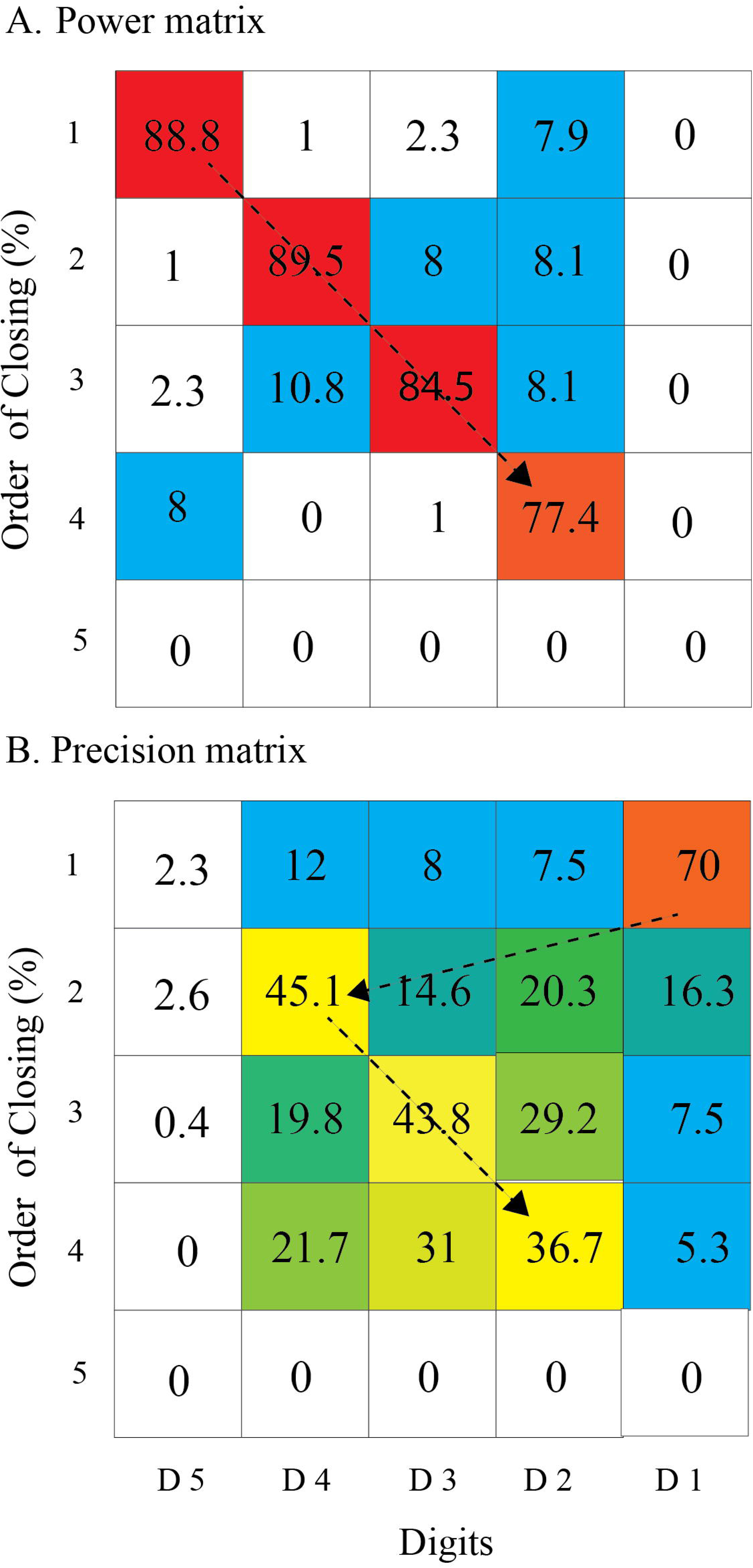
Matrix of digit pad closing to catch. A. Power grip. Matrix indicating the order (percent) of distal digit pad prehension (digit tips approach palm) for reach-to-catch the small ball. Arrows indicate the most frequent closing pattern is digit 5-2, with the reverse pattern 2-5 occurring less frequently. B. Precision grip. Matrix indicating the order (percent) of opposition (distal digit pad closing) for reach-to-catch digit contact with the tennis ball. Arrows indicate the most frequent closing sequence is 1-2-3-4, with other patterns involving these digits occurred frequently. Note: for a power grasp of the small ball the thumb is not involved in grasping and for a precision grasp of the tennis ball, digit 5does not contact the ball but curls under the ball to support it.

As illustrated in Figure 9A (Video 3) principal component and K-Means clustering analysis of power grips identified 4 finger closing clusters comprising 15, 156, 17, and 25 observations respectively. Clusters were:

*Cluster 0*: digit 3 closing at order 1, digit 2 and 4 at order 2, and a mix of digits 3 and 5 at order 3, with digits 2 at order 4 (7% of data points).

*Cluster 1*: digit 5 at order 1 and 2, digit 3 at order 3, and digit 2 at order 4 (73.2% of data points).

*Cluster 2*: digit 2 at order 1, digit 3 at order 2, digits 4 at order 3, and digit 5 at order 4 (8% of data points).

*Cluster 3*: digit 5 at order 1 and 2, digit 3 at order 3, and digit 4 at order 4 (11.7% of data points).

As is illustrated in Figure 9B (Video 4) principal component and K-Means clustering analysis for precision grips identified 4 finger closing clusters comprising 37, 74, 77, and 37 observations respectively. Clusters were:

*Cluster 0*: digits 1 and 2 at order 1, digits 3 at order 2, digits 4 at order 3, and digits 1, 2, and 4 at order 4. (16.4% of data points).

*Cluster 1*: digit 1 at order 1, digit 1 and 4 at order 2, digit 2 and 4 at order 3, and digit 3 at order 4. (32.9% of data points).

*Cluster 2*: digits 1, 3, and 4 at order 1, digit 4 at order 2, digit 3 at order 3, and digit 2 at order 4. (34.2% of data points).

*Cluster 3*: digit 1 at order 1, digit 2 at order 2, digit 3 at order 3, and digit 4 at order 4. (16.4% of data points).

In summary, power grips mainly featured digit closing combinations of digits 5 through 2 (digit 1 was usually not closed on a ball in power grips) and precision grips mainly involved combinations of digits 1 to 4 (digit 5 was not closed on a ball in precision grips of the tennis ball, but supported the ball by curling underneath it. An analysis of hand rotation associated with power grips showed the hand was usually slightly supinated in association with power grip synergy and the hand was slightly pronated with a precision grip synergy. This association of hand rotation and grip synergy has been described by Napier (1956, 1962).

## Discussion

This study examined the hand shaping and grasp types employed by young participants to catch balls of four different sizes and compared these movements with those used to pick up stationary balls or intercept rolling balls. Hand shaping and grasping during catching were sensitive to both intrinsic (size) and extrinsic (movement, location) ball features and differed from those used in picking up and intercepting tasks, as supported by three key observations. First, catching involved seemingly disproportionate hand opening and digit closing up to and including ball contact, although the degree of hand opening remained correlated with target size. Second, grasp type during catching varied with ball size: power grasps were used for the smallest balls, and precision grasps for larger balls. Third, digit closing synergies displayed variability suggesting influence by tactile cues associated with ball contact. Together, these findings align with dual visuomotor channel theory, which proposes that hand movements are mediated by neural pathways sensitive to extrinsic and intrinsic object features (Arbib, 1981; Jeannerod, 1981; Castiello et al., 1992a; Goodale et al., 1994; Hu et al., 2005; Coats et al., 2008; Hoffmann et al., 2010; Karl et al., 2013; Hall et al., 2014).

Previous research has demonstrated that maximum pregrasp aperture (MPA), measured as the distance between the thumb and index finger pads, is proportional to target size (Arbib, 1981; Binkofski et al., 1998; Cavina-Pratesi et al., 2010a, 2010b, 2018; Culham and Valyear, 2006; Fattori et al., 2010; Galletti et al., 2003; Haggard and Wing, 1997; Jeannerod, 1981, 1999; Jeannerod et al., 1994; Karl and Whishaw, 2014; Karl et al., 2012, 2018; Smeets and Brenner, 1999; Vesia et al., 2013; Whishaw et al., 2016) and increases when the target is in motion, such as rolling down an incline (Lacquaniti and Maioli, 1989; Savelsbergh et al., 1991, 1993; Laurent et al., 1994; Button et al., 2002; Mazyn et al., 2006; Tijtgat et al., 2010; Bongers et al., 2012). Consistent with these findings, this study observed similar MPA scaling for stationary and rolling balls. Notably, a much larger increase in MPA was observed when participants caught thrown balls, with the hand appearing fully opened at MPA. Although this disproportionate hand aperture increase has been previously noted for balls in flight (Cesqui et al., 2012), here MPA still scaled with ball size even when the hand was near full opening. Electromagnetic measures confirmed that MPA reflected extension and deviation of all digits, as supported by 3D kinematic measures of finger opposition, digit pad distances, and prehension (distance from digit pads to palm). Thus, while the hand is clearly preshaped when grasping stationary or rolling balls, preshaping appears minimal during catching, as indicated by nearly fully opened MPA.

Previous studies report gradual hand closure before contact during object pickup (Jeannerod, 1981, 1999; Jeannerod et al., 1994; Karl and Whishaw, 2014; Karl et al., 2012, 2018; Smeets and Brenner, 1999a,b). In contrast, this study found that during catching, hand closing at first contact changed only slightly from MPA, with most finger closure occurring post-contact. Thus, both hand opening magnitude and timing of closure differ between catching and reach-to-grasp or reach-to-intercept movements. Consistent with prior literature, participants used precision grasps for static objects and intercepting moving objects. Here, grasp type during catching depended on ball size: smaller balls were caught with power grasps, larger balls with precision grips. Measures of MPA and visual inspection of maximum hand opening did not predict grasp type, aligning with Brenner and Smeets (2018). Nevertheless, smaller balls tended to be contacted the hand’s radial side, while larger balls tended to be contacted near the hand midpoint midpoint suggesting decisions about grasping occurred early during ball flight (Rushton et al., 1999).

A third key finding was that ball contact location on the hand influenced digit synergies for grasping. Frame-by-frame inspection revealsed diverse patterns for power and precision grips, consistent with previous reports for immobile objects (Wong and Whishaw, 2004). Cluster analysis, however, indicated a smaller set of preferred finger sequences: power grips typically involved digits 5-4-3-2 closing, whereas precision grips most often began with digit 1 closing followed by 4, 3, or 2 sometimes one finger preceded the thumb. While Napier (1956, 1962) noted an association between grip type and hand rotation, the variable locations at which the balls were caught from trial to trial prevented a definitive confirmation of this point.

Woodworth’s (1899) formative description of hand movements proposes an initial rapid proximal limb movement followed by refined distal limb adjustments toward a target. This observation formed the basis of dual visuomotor theory: reaching is guided by extrinsic spatial cues, grasping by intrinsic target features (Arbib, 1981, 2016; Jeannerod, 1981; Haggard and Wing, 1997; Jeannerod, 1999). The present findings generally support this view, showing initial hand positioning for interception could be followed by finger closure forming either a power or precision grip depending on ball size. Nevertheless, the control of catching may be still more complex, because somatosensory cues from ball contact contribute to terminal grasp patterning (Johansson and Flanagan, 2009). Possibly for static object acquisition with location certain, vision predominates (Jeannerod, 1981; Castiello et al., 1992b; Chieffi and Gentilucci, 1992; Goodale et al., 1994; Carnahan and McFadyen, 1996; Hu et al., 2005; van de Kamp and Zaal, 2007; Coats et al., 2008; Hoffmann et al., 2010), whereas for moving targets with uncertain location, somatosensation’s role increases. This idea is consitent with findings that reaching into peripheral vision or while blindfolded (Karl and Whishaw, 2012, 2013; Whishaw and Karl, 2014, 2019), tactile feedback from first touch guides hand shaping and grasp selection. Possibly, somatosensory-driven shaping and grasping may involve neural channels parallel to visual ones (Hsndflrert and Ginty, 2021; Magosso, 2010).

In conclusion, participants received no explicit instructions regarding how to pick up, intercept, or catch the balls (Ansuini et al., 2008; Sartori et al., 2011; Valyear et al., 2011), and ball speed and location for catching was not systematically controlled (Chieffi et al., 1992; Carnahan and McFadyen, 1996), preserving response variability and spatial uncertainty. The main findings— effects of ball size on MPA, grip type selection, and somatosensory contributions to finger closing synergies—support the notion of separate visuomotor channels for reach and grasp (Arbib, 1981; Binkofski et al., 1998; Cavina-Pratesi et al., 2010a, 2010b, 2018; Culham and Valyear, 2006; Fattori et al., 2010; Galletti et al., 2003; Haggard and Wing, 1997; Jeannerod, 1981, 1999; Jeannerod et al., 1994; Karl and Whishaw, 2014; Karl et al., 2018; Vesia et al., 2013; Whishaw et al., 2016). Furthermore, the results underscore that the central control of catching integrates visuomotor and somatosensory information from target contact (Johansson and Flanagan, 2009).

## Author contributions

*Conception and design of study: IQW, JRK, AM*

*Provision of study materials and analysis tools: JMK, IQW, MM*

*Performance of experimental methods: JRK, YE, AM, IQW, HR*

*Analysis and interpretation of data: JRK, AM, IQW, YE, HR*

*Conceptualization and writing of manuscript: AM, IQW, JRK*

*Critical revision of manuscript for important intellectual content: AM, IQW, JMK, JBD, AM, MM, YE*

---

**Video 1. Power grip.** A participant makes a power grip catch of the smallest ball using a chase catch strategy in which the hand advances into the ball and the fingers close to clamp it against the palm. (10% normal speed).

**Video 2. Precision grip.** A participant makes a precision grip catch of the largest sized ball using a stop strategy in which the hand remains in relatively the same location as the fingers are closed for purchase. (10% normal speed).

**Video 3. Power grip synergy**. A power grip of the smallest ball in which the digits close in the order 5-4-3-2, in which digit 5 is the pinky. (10% normal speed).

**Video 4. Precision grip synergy**. A precision grip of the third sized ball, a tennis ball, in which the digits close in the order of 1-2-3-4, in which 1 is the thumb. Note that digit 5 does not participate in the precision grip but contributes to providing support on the ball. (10% normal speed).

